# Motion-Corrected Structural Realignment and Quantitative Voxel Targeting in Magnetic Resonance Spectroscopy

**DOI:** 10.1101/2025.07.31.668045

**Authors:** Cassia Low Manting, Atsushi Takahashi, Jyrki Ahveninen, Lilianne R Mujica-Parodi, Eva-Maria Ratai

**Author notes:** Correspondence:* (C.L.M.).

## Abstract

Magnetic resonance spectroscopy (MRS) non-invasively quantifies neurochemical metabolites within a prescribed voxel in the brain, but is highly sensitive to head motion that displaces the voxel from its intended anatomical location, and introduces spectral contamination as well as tissue estimation errors. In particular, voxel displacement into bone tissue severely degrades spectral quality due to lipid contamination, often rendering the data unusable. While promising solutions exist to address motion-related issues, many rely on specialized infrastructure and expertise available only at a limited number of research centers. To address inter-acquisition motion, we combine automatic voxel repositioning using an updated 18 s head scout MRI, acquired immediately before MRS, with realignment of the structural MRI to ensure that tissue correction is based on the updated voxel location. Furthermore to quantify anatomical targeting accuracy, we introduced a voxel–ROI overlap metric that measures the percentage intersection between the prescribed voxel and atlas-defined target regions, enabling objective evaluation of voxel positioning accuracy. Using voxels prescribed across three brain regions (prefrontal cortex, thalamus, left superior temporal gyrus; N = 31), realignment on average increased voxel–ROI overlap by 24%, reduced inter-subject variability in tissue composition by 8.1%, and eliminated bone contamination in all six affected cases. Our computationally efficient method executes in < 35 s per participant and requires no specialized hardware, offering an accessible solution that integrates inter-acquisition motion correction with quantitative assessment to achieve accurate voxel targeting.

## 1. INTRODUCTION

Proton magnetic resonance spectroscopy (MRS) is a noninvasive MRI-based technique that measures the in vivo concentration of neurochemical metabolites such as N-acetylaspartate, creatine, choline, glutamate, GABA, and glutathione (Graaf, 2019). In clinical applications (Bonavita et al., 1999; Burlina et al., 2000; Coelho et al., 2025; Dager et al., 2008; Öz et al., 2014; Ratai et al., 2013, 2018; Robertson et al., 2016), it enables the detection of metabolic abnormalities that may not be visible on structural MRI, revealing region-specific neurochemical alterations linked to disease pathology or treatment response. For example, MRS has been applied in neurodegenerative diseases (e.g., Alzheimer’s, Parkinson’s), tumors, and psychiatric conditions including schizophrenia and depression. A critical challenge in MRS is its sensitivity to participant motion, which can drastically degrade data quality and reliability of metabolite quantification (Saleh et al., 2020). Since MRS acquires spectra from a predefined voxel (or voxels in chemical shift imaging) localized to a specific brain region, participant motion displaces the voxel from its original intended location, resulting in metabolite measurements originating partly or entirely from the wrong location. This problem is exacerbated when targeting small regions such as the hippocampus, where motion can cause the voxel to miss the region completely. If undetected, which is often the case without obvious spectral artefacts, the acquired data would then reflect a different anatomical region than intended, compromising reproducibility. For superficial regions, even slight motion may shift the voxel into adjacent bone or scalp tissue, introducing lipid contamination that can dominate the spectra, obscure neurometabolite peaks, and in severe cases render the data unusable. Furthermore, voxel displacement toward air–tissue interfaces increases susceptibility to B_0_ field distortions, which can alter resonance frequencies and disrupt techniques such as water suppression and J-difference editing (Deelchand et al., 2019; Keating & Ernst, 2012). Even when the voxel remains entirely within brain tissue and spectral quality appears acceptable, motion-induced changes in the sampled proportions of grey matter, white matter, and cerebrospinal fluid (CSF) can introduce errors in partial volume correction and metabolite concentration estimates (Archibald et al., 2026). Because these errors frequently occur without obvious spectral artefacts, they may remain undetected while reducing the accuracy and reproducibility of metabolite quantification. To make matters worse, the above problems are magnified at ultra-high field strengths such as 7T, which is gaining popularity for its improved signal-to-noise ratio and spectral resolution, albeit at the cost of increased motion sensitivity. At 7T, the larger chemical shift displacement error significantly increases the risk of lipid contamination from motion, while steeper magnetic susceptibility gradients intensify B_0_ field distortion—particularly near air-tissue and bone-brain interfaces—thereby exacerbating motion-related spectral line broadening and signal degradation compared to 3T (Godlewska et al., 2017).

MRS studies often involve patient populations (e.g. Parkinson’s, multiple sclerosis, dementia) and children who may struggle to remain still during scanning, as well as long scan durations that increase the likelihood of motion (Fox et al., 2011; Saleh et al., 2020). Furthermore, it is not uncommon to acquire the MRS after a substantial period from the initial structural MR scan, for example, when targeting multiple voxel locations, or when fMRI sequences are interleaved in between. This markedly increases the risk of inter-acquisition participant movement, resulting in voxel misplacement relative to the intended location. A plausible solution might be to reacquire the structural MRI and re-plan the MRS voxel in the correct location right before beginning the MRS acquisition. In practice, however, high-resolution structural MR scans require several minutes each—almost comparable to the MRS acquisition itself—and repeating them for every voxel would significantly extend the total scan time, which is both inefficient and often infeasible due to resource constraints. Over the years, researchers have developed several approaches to implement motion correction in MRS. Among the most effective are real-time motion correction methods (Adanyeguh et al., 2024; Andrews-Shigaki et al., 2011; Bogner et al., 2014; Deelchand et al., 2019; Felblinger et al., 1998; Hess et al., 2011; Keating et al., 2010; Keating & Ernst, 2012; Saleh et al., 2016; Wilson, 2019; Zaitsev et al., 2010) that use external motion tracking devices or MRI-based navigator sequences to detect and correct for head motion during acquisition (see Saleh et al. (Saleh et al., 2020) for an in-depth review). However, implementing these solutions is complex – typically requiring specialized hardware, software, or expertise that are not readily accessible to standard users. Because of this complexity, real-time motion correction is currently limited to only a few specialized research centers.

To mitigate voxel misplacement arising from inter-acquisition participant movement, we propose to first acquire an 18 s head scout before each MRS voxel acquisition and reposition the voxel based on the updated scout using AutoAlign (Dou et al., 2015; van der Kouwe et al., 2005) (available in most modern MRI scanners) to retain its intended location. While previous studies have demonstrated methods for improving voxel placement consistency (Ratai et al., 2008), updating the voxel position alone is insufficient, as tissue segmentation relies on the higher-resolution T1-weighted anatomical image that is no longer aligned with the repositioned voxel. Segmentation is an essential step in MRS analysis that computes grey matter, white matter, and CSF volumes within the MRS voxel to enable accurate metabolite quantification by correcting for tissue-specific contributions. Recent work has further demonstrated that tissue segmentation strategies themselves can influence metabolite quantification, highlighting the critical role of tissue segmentation in MRS analysis (Archibald et al., 2026). Hence to ensure that segmentation and tissue correction were performed in motion-corrected space, we realigned the original T1-weighted image and tissue probability maps to the updated scout position using a computationally efficient algorithm (< 30s runtime). This process involved first extracting the parameters that transform the pre-to post-motion head scout, then applying these parameters to the T1-weighted image and tissue probability maps to spatially align them with the post-motion head scout (See **Methods – D. Affine Realignment of T1-weighted Image and tissue probability maps**). This supports more accurate metabolite estimation that accounts for motion between acquisitions—effectively replicating the benefit of acquiring a new T1-weighted image but with over a ten-fold reduction in acquisition time. Such an approach is especially beneficial in motion-prone settings, offering a quick and practical fix that is highly accessible to new users while real-time motion correction technologies continue to develop towards widespread adoption. Similarly, longitudinal studies require consistent anatomical sampling across repeated imaging sessions, making reliable voxel positioning important not only within a scanning session but also across time. Furthermore, despite the critical importance of accurate voxel placement in MRS, quantitative metrics for objectively evaluating anatomical accuracy of voxel positioning are rarely adopted or reported in practice. Assessment of voxel placement accuracy is generally limited to visual inspection of a representative example, rather than quantitatively evaluated across all participants. To address this limitation, we developed a voxel–ROI overlap metric that quantifies the volumetric intersection between the prescribed voxel and atlas-defined anatomical regions. This metric enables objective evaluation of voxel displacement, post motion-correction recovery accuracy, and inter-subject consistency, and longitudinal consistency of anatomical sampling. Fig. 1. illustrates an overview of the proposed integrated framework, showing how the realignment and voxel-ROI overlap algorithms fit within the MRS analysis pipeline. In this paper, we demonstrate the success of our structural realignment approach by showing that displaced voxels were restored to their original intended locations, improving overlap with the target region-of-interest, eliminating bone contamination, and reducing inter-subject variability in grey matter, white matter, and CSF volumes within the voxel.

**Fig. 1.**
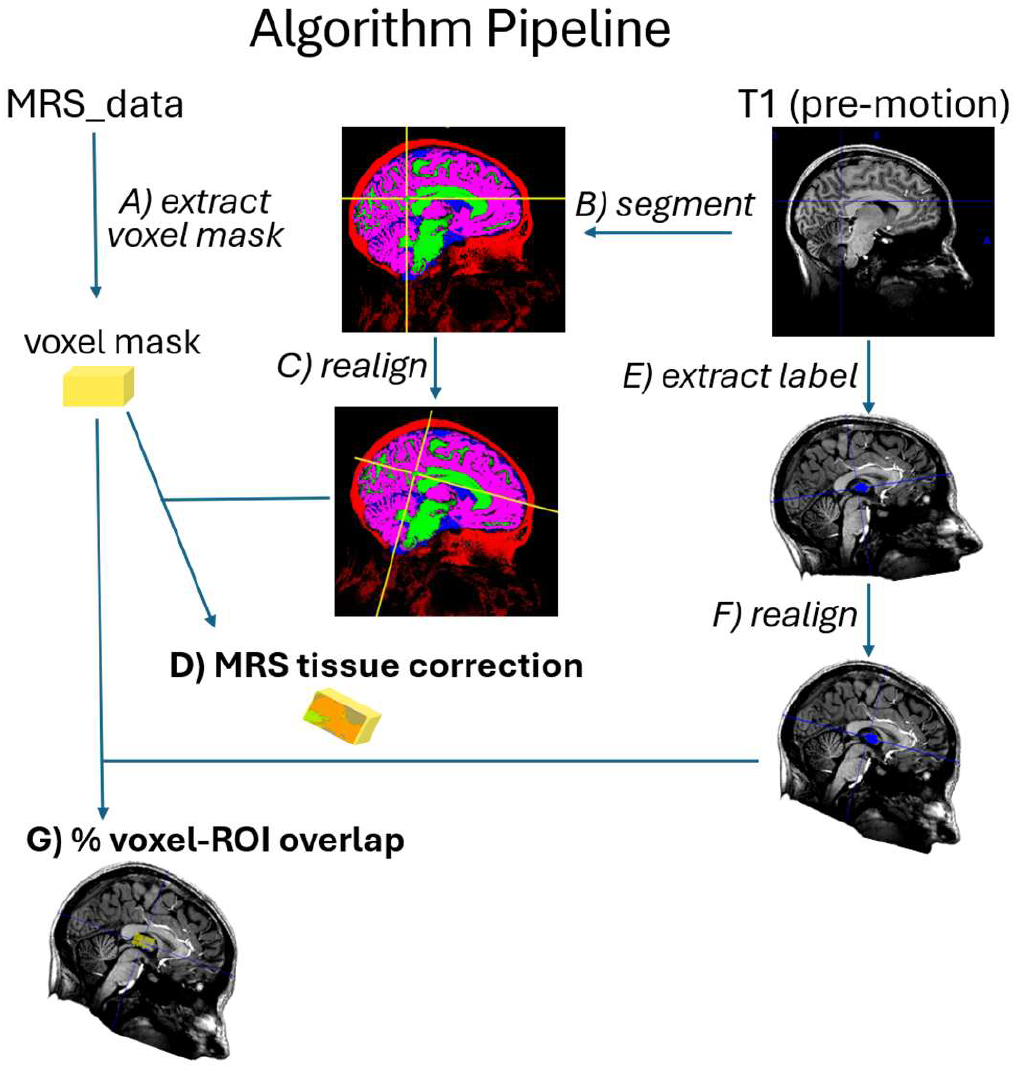
Overview of algorithm pipeline, including both realignment and voxel-ROI overlap. A) The voxel mask was extracted from the MRS acquisition data file using FSL-MRS’s *spec2nii* (Clarke et al., 2021). B) The pre-motion T1 MRI was segmented to produce grey matter (magenta), white matter (green), CSF (blue), and bone (red) tissue probability maps using a modified version of *Gannet_segment*. C) The tissue probability maps were then realigned to the updated post-motion position based on pre- and post- motion headscouts using Freesurfer’s *mri_robust_register* and *mri_vol2vol*. D) The extracted voxel mask and realigned tissue probability maps were used for accurate MRS tissue correction. E) Anatomical labels were extracted using Freesurfer’s *recon-all* (Thalamus is labelled in blue in the above example). F) The anatomical labels were realigned to the updated post-motion position as in C). G) The % voxel-ROI overlap was then computed as the percentage volume of voxel mask (ýellow) that intersects with the anatomical label (blue).

## 2. Methods

### 2.1. Participants

Data were collected from 31 healthy participants (17 female), aged 22 to 79 years (mean age = 48.5 ± 14.9 years), who were recruited from the Boston metropolitan area via online and print advertisements. The present study re-analyses a subset of data previously reported by Hone-Blanchet (Hone-Blanchet et al., 2023) and Nieuwenhuizen et al. (van Nieuwenhuizen et al., 2025) to focus on motion-related effects. The study was registered on ClinicalTrials.gov (NCT04106882) and approved by the institutional review boards of Massachusetts General Hospital (2015P000652) and Stony Brook University (IRB2019-00208). Written informed consent was obtained from all participants prior to participation.

### 2.2 Structural Magnetic Resonance Imaging (MRI)

Structural MRI data were acquired on a 7 Tesla MR scanner (Siemens Healthineers, Erlangen, Germany) equipped with a 32-channel head coil developed in-house (Athinoula A. Martinos Center for Biomedical Imaging, Massachusetts General Hospital, USA). High-resolution T1-weighted images were collected using a multi-echo magnetization-prepared rapid gradient echo (MEMPRAGE) sequence at 1 mm isotropic resolution (TA = 6 min 3 s, TE = 1.61, 3.47, 5.33, and 7.19 ms; TR = 2530 ms; flip angle = 7°; FOV = 256 x 256 mm; 176 sagittal slices; GRAPPA, R = 2, phase-encoding direction, 32 ACS lines). For each participant, two 18 s low-resolution 3D AutoAlign head scouts were acquired at 1.6 mm isotropic resolution (TA = 17 s; TE = 1.53 ms; TR = 4.00 ms; flip angle = 16°; FOV = 260 × 260 mm; 128 sagittal slices; GRAPPA, R = 3, phase-encoding direction, 24 ACS lines) approximately 77 minutes apart, during which other data (e.g. fMRI) outside the scope of this paper was acquired. Furthermore, participants were not given any specific instructions to move. To evaluate the current framework, voxel repositioning was retrospectively simulated using the head scouts to reproduce the scanner’s automated voxel repositioning function that aligns the prescribed MRS voxel to the same location based on anatomical landmarks.

### 2.3 Voxel Positions and Dimensions

Three voxels were placed following common neuroanatomical guidelines and spectroscopy dimensions (reported in LR-AP-SI format): a i) 20 × 20 × 20 mm^3^ voxel in the prefrontal cortex (PFC), centered between the rostral anterior cingulate cortex and the frontal pole, ii) 34 × 21 × 14 mm^3^ voxel in the thalamus (Th), iii) 14 × 34 × 21 mm^3^ voxel in the left superior temporal gyrus (LSTG), centered at Heschl’s gyrus and rotated to align maximally along the superior temporal gyrus. The last two dimensions were selected to reflect the larger voxel sizes for better signal-to-noise required by J-edited MRS sequences. The axial, sagittal, and coronal views of these voxel positions are illustrated in Fig. 3. Automated voxel positioning, such as Siemen’s AutoAlign (Dou et al., 2015; van der Kouwe et al., 2005) or Philip’s SmartExam (Young et al., 2006), should be used to automatically recenter voxels according to updated head scout acquisitions, maintaining voxel placement in the intended location.

### 2.4 Voxel segmentation and tissue composition

Pre-motion anatomical MRIs were segmented into grey matter (GM), white matter (WM), cerebrospinal fluid (CSF), and bone partial volume maps using the unified tissue segmentation algorithm in SPM12 (https://www.fil.ion.ucl.ac.uk/spm/) with default settings. Tissue fractions, converted to percentages, were then computed by sampling the corresponding probabilistic maps inside the binary MRS voxel mask, summing each tissue’s probabilities, and dividing by the total GM + WM + CSF + bone probability. These steps were implemented using Gannet_Segment5.m, modified from the default Gannet (Edden et al., 2014) segmentation function to incorporate bone segmentation. The algorithm describes the generation of tissue-specific probabilistic maps (GM, WM, CSF, and bone), followed by multiplication with the binary voxel mask to obtain voxel-restricted tissue maps. Tissue fractions were computed by summing these voxel-wise probabilities and normalizing by the total tissue probability within the voxel, thereby accounting for partial volume contributions across all tissue classes. The source code for Gannet_Segment5.m is publicly available at https://github.com/cassialmt/MRSmotion_publication. As subject motion may increase or decrease individual GM, WM, and CSF tissue fractions depending on the direction of voxel displacement, realignment performance was evaluated by restoration toward the original voxel composition and reduction in inter-subject variability, reflecting improved consistency of anatomical sampling, rather than systematic changes in group mean tissue fractions.

### 2.5 Affine Realignment of T1-weighted Image and tissue probability maps

For each participant, two head scout images were acquired ∼77 min apart and affinely aligned by estimating the rotation and translation needed to overlay the images with minimal intensity differences between corresponding voxels. These steps can be mathematically described as follows (Fig. 2A):

**Fig. 2.**
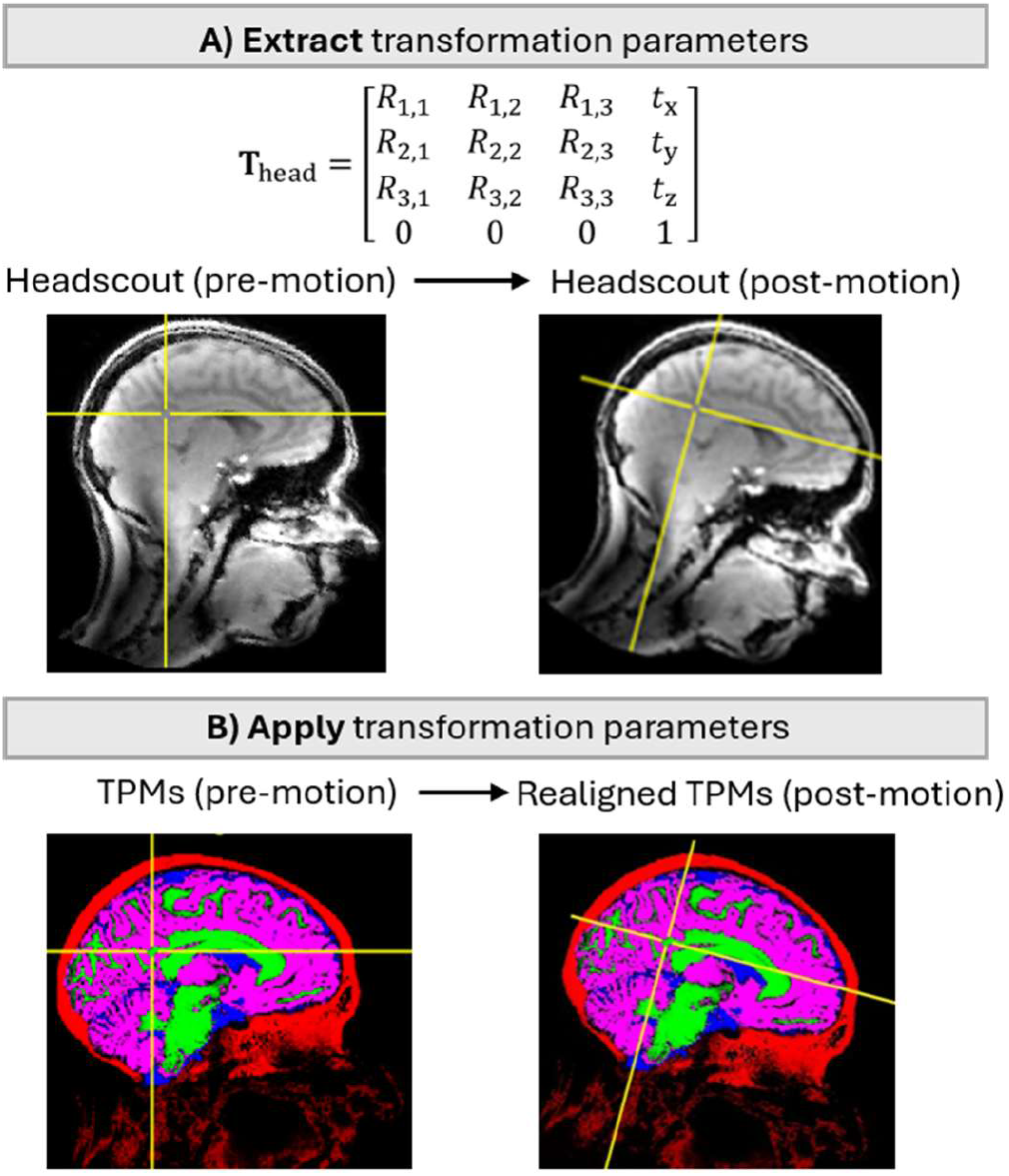
Detailed implementation of the structural realignment procedure. A) For each participant, pre- and post- motion head scout scans were affinely aligned to estimate the transformation parameters **T**_head_. B) The transformation was then applied to realign the pre-motion tissue probability maps (TPMs) to match the post-motion position, avoiding the need to reacquire a post-motion T1 image.

Let *H*_pre_ and *H*_post_ denote the head scout images acquired before and after motion respectively. The affine transformation matrix *T*_head_ aligning *H*_pre_ to *H*_post_ was estimated by minimizing the voxel-wise squared intensity differences:

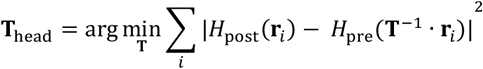

where *r*_i_ indexes over voxel coordinates in the image space. *T*_head_ *∈* ℝ^4^ is the 4×4 affine transformation matrix including rotation and translation, estimated by FreeSurfer’s (Fischl, 2012; Reuter et al., 2010) *mri_robust_register* with automatic saturation estimation:

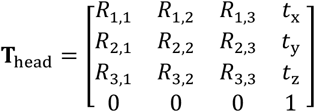

This transformation was then applied to realign the original T1-weighted image acquired immediately after the first head scout, and the tissue probability maps obtained from segmentation, to the post-motion head scout position to produce corresponding post-motion equivalents (Fig. 2B). This operation was implemented using the FreeSurfer (Fischl, 2012) command *mri_vol2vol* with nearest-neighbor interpolation. Nearest-neighbour interpolation was selected to preserve the original image intensities and native tissue contrast while avoiding artificial smoothing prior to segmentation and anatomical labelling. As the applied transformations were purely affine and of limited magnitude, the potential for introducing substantial staircase-like artifacts at tissue boundaries was relatively low. Visual quality control confirmed that cortical and subcortical boundaries remained anatomically plausible and that no systematic distortions were introduced by the selected interpolation method. The total runtime for both commands was ∼35 s per participant. FreeSurfer is an open-source software package available at https://surfer.nmr.mgh.harvard.edu.

### 2.6 Head motion

Head motion was quantified using the 4×4 affine transformation matrix obtained previously (see previous section). Net translation was calculated as the Euclidean norm of the translation vector *∥*t*∥* (last column of the first three rows), where:

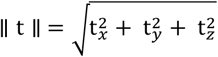

Net rotation was determined by extracting Euler angles (θ_x_, θ_y_, θ_z_) from the upper-left 3×3 rotation submatrix *R* via the formula:

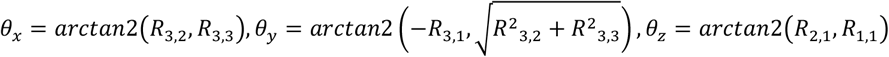

The overall rotation magnitude *∥*θ*∥* was defined as the Euclidean norm of (θ_X_, θ_y_, θ_z_), where:

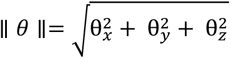

### 2.7 Quantitative voxel-ROI overlap metric

To objectively evaluate anatomical targeting accuracy, we defined a voxel-ROI overlap metric as the percentage of voxel volume *V* intersecting with the target anatomical region *R* (Fig. 3):

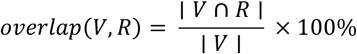

where *∣ V* ∩ *R ∣* denotes the volume of intersection between the voxel and ROI, and *∣ V ∣* denotes total voxel volume. The computational implementation used for individual-level voxel mask and anatomical label extraction, as well as voxel-ROI overlap calculation is publicly available at https://github.com/cassialmt/MRSmotion_publication.

ROIs were as defined on the pre-motion T1-weighted MRI by Freesurfer’s subcortical atlas (Fischl et al., 2002) (Th) and the Deskian-Killiany atlas (Desikan et al., 2006) (PFC/LSTG). PFC encompasses the bihemispheric medial orbital frontal and superior frontal labels, LSTG corresponds to the left Heschl’s gyrus and superior temporal gyrus labels, and Th corresponds to the thalamus label. For computation of the realigned voxel-ROI overlap, anatomical labels were transformed using *T*_head_, estimated from aligning the pre-and post-motion head scouts in Section 2.5. Mean differences in voxel–ROI overlap across participants before and after realignment were assessed using two-tailed paired *t*-tests, with effect sizes reported as Cohen’s *d* for paired samples (mean difference divided by standard deviation).

**Fig. 3.**
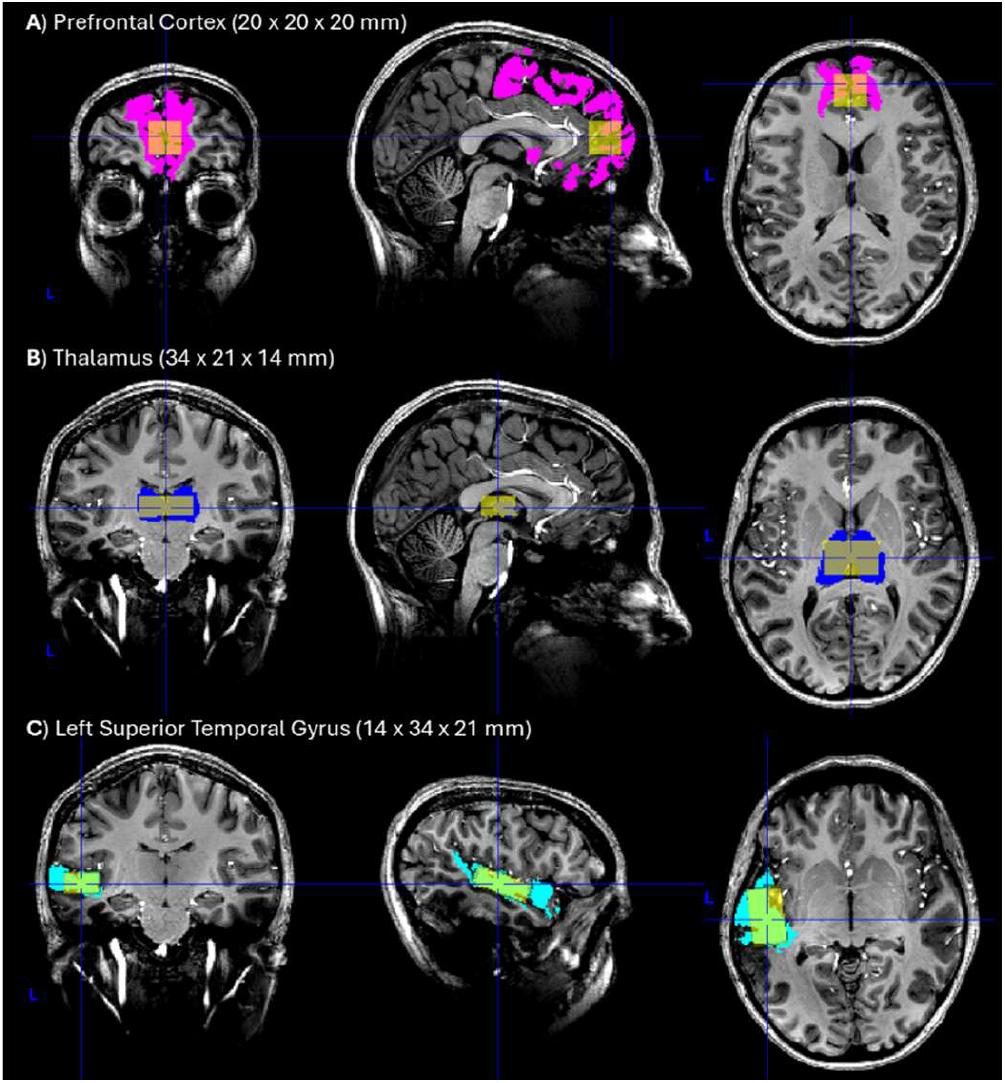
Voxel-ROI overlap between target voxel and atlas-defined anatomical region. Three voxels (yellow) were placed following common spectroscopy dimensions (in parentheses), in the **(A)** prefrontal cortex (magenta), centered between the rostral anterior cingulate cortex and the frontal pole, **(B)** thalamus (blue), and **(C)** left superior temporal gyrus (cyan), centered at Heschl’s gyrus and rotated for maximal overlap. The voxel-ROI overlap is calculated as the percentage of voxel volume intersecting with the target anatomical region.

## 3. Results

The proposed realignment method was evaluated using a dataset (N = 31) containing, for each participant, two low-resolution head scouts acquired ∼77 minutes apart and a T1-weighted structural MRI obtained immediately after the first head scout. Analyses were performed for voxels placed in the prefrontal cortex (PFC), thalamus (Th), and left superior temporal gyrus (LSTG) (see Fig. 3 for exact voxel locations and dimensions). As an independent validation, the realigned T1-weighted image was compared against a post-motion T1-weighted MRI acquired immediately after the second head scout. The two images showed close agreement, with a mean translation of 1.31 ± 0.84 (range: 0.32 - 4.02) mm and mean rotation of 0.020 ± 0.015° (range: 0.002 - 0.069°). Because the post-motion T1-weighted image was acquired after the second head scout, these residual differences likely reflect head motion occurring between the two acquisitions rather than inaccuracies of the proposed realignment itself.

### 3.1 Voxel-ROI overlap

After ∼77 minutes, participant motion (sigma translation = 6.61 ± 4.27 (range: 1.24 - 22.68) mm; mean rotation = 2.85 ± 1.42° (range: 0.89 – 7.18°).; Fig. 4A) substantially displaced the original voxel positions, resulting in reduced overlap with the intended ROIs. Fig. 4B provides a conceptual illustration of how head motion can displace prescribed voxels relative to their target anatomical location. The proposed realignment method seeks to correct this displacement by restoring voxels to their intended position. The voxel-ROI overlap was computed to evaluate the anatomical accuracy of voxel placement at each target ROI. It is calculated as the percentage of voxel volume intersecting with the target ROI (Fig. 4C), as defined by standard anatomical atlases (see **Methods**). Following realignment, voxel positions were successfully restored to their original locations, recovering the initial voxel-ROI overlap in all participants (Fig. 4C). Across participants, voxel–ROI overlap increased significantly from 39.0 ± 8.7% to 57.4 ± 7.2% in PFC (mean = 18.4%, 95% CI [15.1, 21.6], *t*(30) = 11.59, *p* < 0.0001, *d* = 2.08), 55.6 ± 18.5% to 81.4 ± 4.8% in Th (mean = 25.8%, 95% CI [19.2, 32.4], *t*(30) = 8.02, *p* < 0.0001, *d* = 1.44), and 39.2 ± 17.1% to 67.9 ± 8.0% in LSTG (mean = 28.6%, 95% CI [22.8, 34.5], *t*(30) = 9.99, *p* < 0.0001, *d* = 1.79). Increase in voxel–ROI overlap was observed across all regions for every participant, except one individual who showed a slight 1.2 % decrease for the PFC. Furthermore, the reduction in inter-subject variance (equivalent to standard deviation squared) of voxel-ROI overlap indicates improved spatial consistency of voxel placements across participants.

**Fig. 4.**
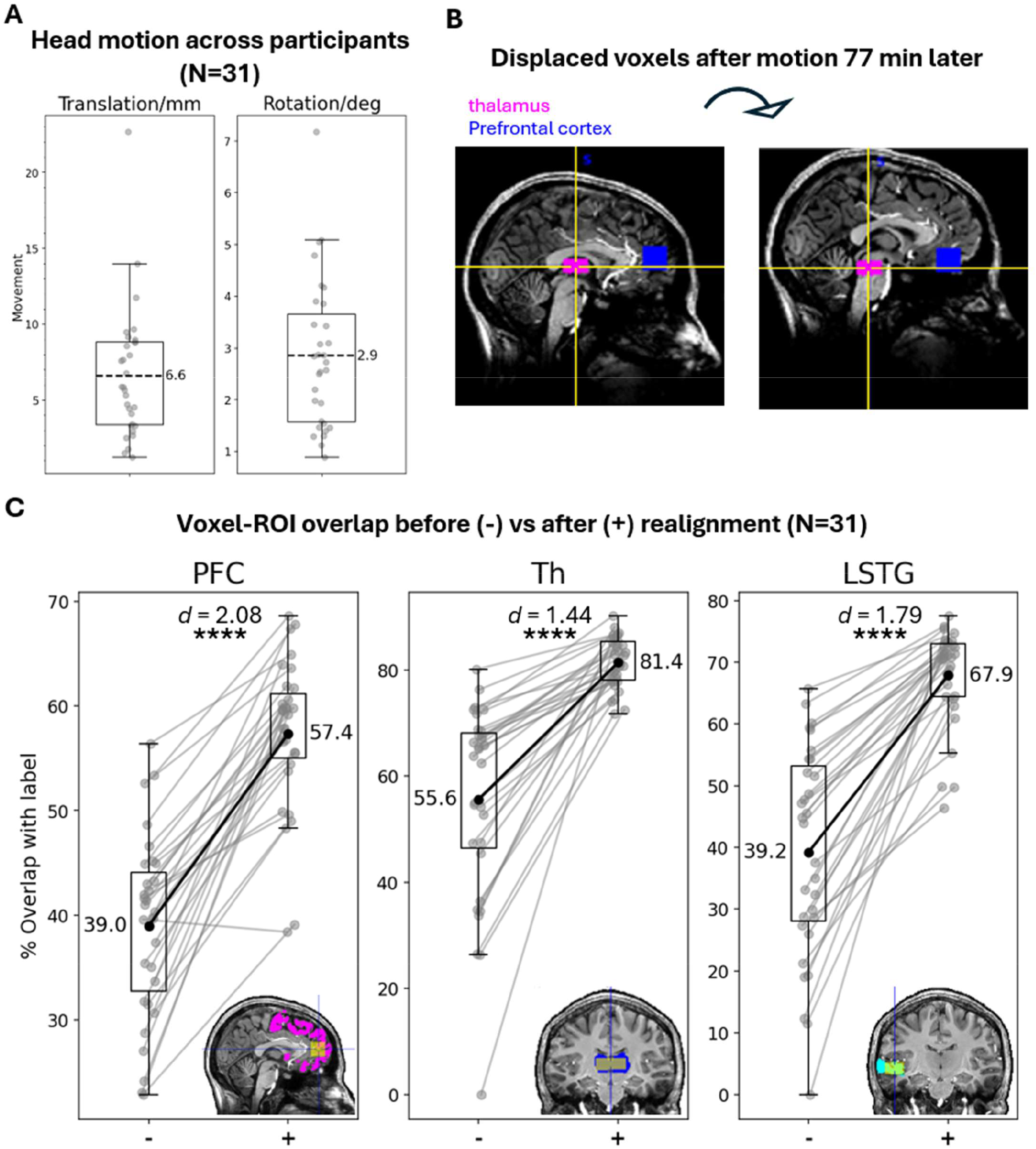
Realignment corrects inter-acquisition motion and restores voxel-ROI overlap. **(A)** Head motion between head scout acquisitions across participants (individual markers; N = 31). For each individual, translation (left) and rotation (right) were computed from two MR head scout images acquired ∼77 minutes apart, resulting in a mean translation and rotation of 6.61 mm and 2.85° respectively (dotted horizontal lines). **(B)** Illustrative example demonstrating how head movement displaced prescribed voxels in the thalamus (magenta) and prefrontal cortex (blue) from their intended locations in a single participant. This figure is intended for visualization only and is not part of the realignment methodology or analysis. **(C)** Realignment improved voxel–ROI overlap across all regions. Mean overlap (displayed) increased significantly from 39.0 to 57.4% in PFC, 55.6 to 81.4 % in Th, and 39.2 to 67.9% in LSTG (\*\*\*\**p* < 0.0001). Every participant, except one, showed increased overlap in all regions. Furthermore, inter-subject variability of the overlap decreased, reflecting more consistent voxel placements across participants. Box plots outline the 25th – 75th percentiles, with whiskers extending to non-outlier data points. The voxel-ROI overlap was computed as the percentage volume of voxel (yellow) intersecting with the target ROI (colored region), as defined by standard atlases. The lower-right inset in each column illustrates the overlap.

### 3.2 Voxel tissue composition

We further analyzed changes in voxel tissue composition, namely the percentage volume of grey matter (GM), white matter (WM), cerebrospinal fluid (CSF), and bone, before and after realignment. In all six cases where the voxel included > 1% bone volume before realignment, realignment eliminated bone contamination completely (Fig. 5A and B). This is critical as even minimal lipid contamination arising from bone marrow can render an MRS spectra completely unusable due to excessive noise (Saleh et al., 2020). Following alignment, grey and white matter volumes returned to original levels, which were typically higher or similar—except in a few cases where voxels were coincidentally moved to regions containing more grey and white matter (Fig. 5C). The mean percentage grey and white matter volume increased from 82.6 ± 7.3% to 82.7 ± 7.1% in PFC, 81.2 ± 12.1% to 86.7 ± 4.2% in Th, and 84.0 ± 9.6% to 85.7 ± 5.3% in LSTG. CSF volumes similarly reverted to original values after alignment (Fig. 5D). The mean percentage CSF volume changed from 16.5 ± 7.2% to 17.1 ± 7.0% in PFC, 18.7 ± 12.1% to 13.2 ± 4.2% in Th, and 13.3 ± 7.0% to 13.3 ± 5.4% in LSTG. Importantly, these changes were generally accompanied by reduced inter-subject variability in voxel GM, WM, and CSF compositions. Such consistency in voxel tissue composition is crucial for accurate MRS metabolite quantification as changes in grey matter, white matter, and CSF fractions can systematically bias metabolite estimates even when spectral quality appears acceptable. To assess the practical impact of reduced inter-subject tissue variability, we simulated CSF-corrected metabolite estimates before and after structural realignment (Supplementary Material S1). Structural realignment reduced inter-subject variability, decreasing the mean standard deviation by 80.4% in the Tha, 32.9% in the LSTG, and 1.5% in the PFC, indicating improved consistency and reproducibility of metabolite quantification. Together with increased voxel-ROI overlap, these findings indicate more consistent voxel placement within the intended anatomical regions following realignment.

**Fig. 5.**
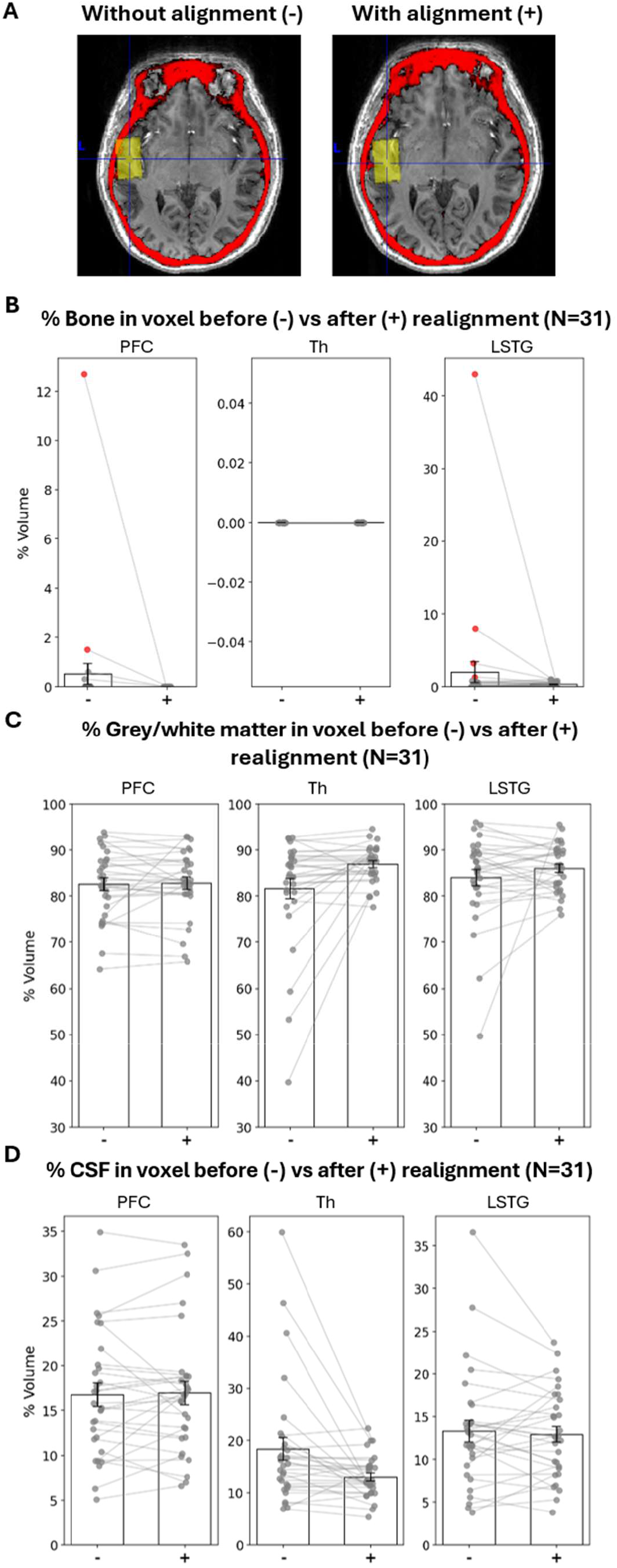
Effects of realignment on voxel tissue composition. (A) Voxel bone contamination before and after realignment. The above example shows the LSTG voxel (yellow) intersecting with skull bone (red) at the left-frontal edge (left panel), which was corrected following realignment (right panel). (B) Elimination of bone contamination. Realignment eliminated bone contamination entirely in all six affected cases (red), eliminating the risk of bone lipid contamination. (C) Grey and white matter volumes were restored to original levels following realignment. Group mean (N=31) percentages increased from 82.6 to 82.7% in PFC, 81.2 to 86.7% in Th, and 84.0 to 85.7% in LSTG. (D) Cerebrospinal fluid (CSF) volumes were similarly restored from 16.5 to 17.1 % in PFC, 18.7 to 13.2 % in Th, and 13.4 to 13.3 % in LSTG. Importantly, the accompanying general reduction in inter-subject variability reflects more consistent voxel tissue composition, which underpins reliable metabolite quantification. Bar plots show means ± standard error of the mean (SEM). LSTG, left superior temporal gyrus; PFC, prefrontal cortex; Th, thalamus;

## 4. Discussion

Effective motion mitigation is essential for accurate MRS quantification. Addressing motion in MRS is a multi-faceted challenge—ranging from prevention (e.g., restraints or shorter scans), to real-time detection (using external trackers or navigators), followed by prospective (Adanyeguh et al., 2024; Andrews-Shigaki et al., 2011; Bogner et al., 2014; Deelchand et al., 2019; Hess et al., 2011; Keating et al., 2010; Keating & Ernst, 2012; Saleh et al., 2016; Zaitsev et al., 2010) or retrospective correction (Felblinger et al., 1998; Ratai et al., 2008; Saleh et al., 2020; Wilson, 2019). To address voxel misplacement caused by motion between acquisitions, we implemented a fast, scout-based realignment strategy using automatic voxel repositioning and post hoc transformation of the T1-weighted MRI as well as tissue probability maps. Unlike previous methods, our strategy requires no additional equipment, lengthy pulse sequences, or specialized in-house expertise, relying primarily on combining various open-sourced tools with customized algorithms. It also avoids estimation-based corrections, eliminating biases introduced by model assumptions and extrapolation. Our results demonstrate improved voxel targeting, elimination of bone contamination, and restoration of consistent tissue composition, making this a practical solution for motion-prone studies, particularly when targeting regions susceptible to artefact contamination that are near bone or air spaces (e.g. cortical surface, sinuses). The benefits of accurate voxel realignment extend beyond preventing catastrophic spectral failure caused by lipid contamination, which often produces obvious artefacts that lead to dataset rejection. More subtle motion-induced changes in voxel tissue composition, particularly CSF fraction, may instead leave conventional spectral quality metrics largely unaffected while systematically biasing metabolite quantification. Consequently, accurate anatomical targeting is essential not only for preventing unusable acquisitions but also for minimizing these otherwise silent sources of quantitative error. This method can also be used across different scan sessions within the same participant to maintain highly consistent voxel placement while obviating the need to reacquire another T1 image. An important limitation of the present study is that it evaluates anatomical rather than spectroscopy-derived outcomes such as metabolite estimates, Cramer-Rao Lower Bound values, linewidth, and signal-to-noise ratio. Nevertheless, because accurate voxel placement and tissue composition are fundamental determinants of metabolite quantification, the present work demonstrates restoration of these anatomical prerequisites that support more accurate metabolite estimation. It should be noted that while a dataset with substantial inter-scan motion was intentionally selected to evaluate the robustness of our realignment method under extreme conditions, the original experiment included an additional T1 acquisition immediately prior to the later MRS as a precaution against motion. While the proposed realignment method is straightforward and easy to implement, it does not correct for dynamic B_0_ fluctuations, motion occurring after the second head scout acquisition, or intra-acquisition motion during the MRS acquisition itself, which can be especially problematic in motion-prone pediatric and patient populations (Fox et al., 2011; Hess et al., 2014). Nevertheless, it offers a quick and highly accessible strategy against motion that may serve as a valuable addition to MRS acquisition protocols where accurate anatomical targeting is essential. Rather than relying on the conventional assumption of negligible participant motion, this approach incurs only an additional 18 s of acquisition time while offering greater security and confidence in voxel placement accuracy.

In addition to correcting segmentation space, the present work introduced a voxel–ROI overlap metric to provide quantitative evaluation of anatomical targeting accuracy in MRS. While the present work focuses on voxel overlap with the target region, interpretation of MRS results can benefit from consideration of the complete anatomical composition of the voxel, including contributions from tissue surrounding the target ROI. In practice, a substantial fraction of the voxel signal may arise from tissue surrounding the ROI, particularly in small or irregularly shaped targets such as the hippocampus. These surrounding contributions may differ systematically across studies due to variations in voxel size, orientation, and placement conventions. Systematic reporting of full voxel composition, including grey matter, white matter, CSF, target and non-target regions, may therefore enhance transparency, cross-study comparability, and confidence in shared MRS findings. Beyond descriptive reporting, voxel composition metrics may be incorporated directly into statistical modeling. Inter-individual variability in anatomical overlap introduces systematic differences in tissue sampling that may confound metabolite comparisons. Including anatomical composition measures as covariates in group-level analyses enables correction for these differences, thereby improving interpretability of metabolite concentration estimates. This approach may be particularly relevant in longitudinal studies or multi-site datasets where subtle placement variability is unavoidable. Integrating voxel composition metrics into statistical models represents an important step toward more robust, replicable, and anatomically informed MRS analysis pipelines. The voxel–ROI overlap metric may also be applied in reverse: rather than solely evaluating anatomical accuracy post hoc, overlap metrics can also be applied prospectively to guide voxel design. By computing expected overlap between candidate voxels and atlas-defined ROIs, researchers can optimize voxel geometry and placement prior to acquisition. This approach may be particularly valuable in small structures (e.g., hippocampus, brainstem nuclei), where trade-offs between signal-to-noise ratio and anatomical specificity are critical. Thus, voxel–ROI overlap analysis can function not only as a validation tool but also to quantitatively inform MRS protocol design and development. In the present study, the voxel–ROI overlap metric was normalized by voxel volume to quantify the contribution of the target ROI to the acquired MRS signal. Alternatively, the metric can be normalized by ROI volume to quantify the proportion of the anatomical ROI covered by the MRS voxel when that is the objective. Finally, the developed voxel-ROI overlap metric may be extended to other quantitative MRI modalities beyond spectroscopy, such as functional MRI, and longitudinal studies requiring consistent ROI sampling.

## Data and Code Availability

All code is publicly available at https://github.com/cassialmt/MRSmotion_publication.

## Author Contributions

C.M.: Conceptualization, Data curation, Formal analysis, Funding acquisition, Investigation, Methodology, Project administration, Software, Validation, Visualization, Writing – original draft, Writing – review & editing. E.R.: Conceptualization, Project administration, Resources, Supervision, Validation, Writing – review & editing. L.M.: Data curation, Funding acquisition, Investigation, Project administration. J.A.: Funding acquisition, Resources, Supervision, Validation, Writing – review & editing. A.T.: Investigation, Methodology, Resources.

## Funding

This research was funded by the W.M.Keck Foundation (to L.M.) and the NSF Brain Research through Advancing Innovative Neurotechnologies Initiative (NCS-FR 1926 781 to L.M.). This study was also supported by The Swedish Foundation for International Cooperation in Research and Higher Education (PD2023-9291 to C.M.).

## Declaration of Competing Interests

The authors declare no competing interests.

## Acknowledgements

We thank Antoine Hone-Blanchet for collecting the data used in this study.

## Notes

### Competing Interest Statement

The authors have declared no competing interest.

### Summary of Updates

co-author requested changes to his description at end of manuscript

