## Supplementary Material for "Motion-Corrected Structural Realignment and Quantitative Voxel Targeting in Magnetic Resonance Spectroscopy"

**S1. Effect of Reduced CSF Variability on Tissue-Corrected Metabolite Estimates**

Changes in cerebrospinal fluid (CSF) fraction within an MRS voxel considerably influence tissue-corrected metabolite estimates because CSF is assumed to contain negligible metabolite signal. To illustrate the practical implications of the observed reduction in voxel tissue variability following structural realignment, we performed a simple simulation based on standard CSF correction given by:

$$C_{\mathrm{corr}} = \frac{C_{\mathrm{meas}}}{1 - f_{\mathrm{CSF}}}$$

$$= \frac{1}{1 - f_{\mathrm{CSF}}}$$

when $C_{\mathrm{meas}} = 1$ and

- $C_{\mathrm{corr}}$ is the tissue-corrected metabolite concentration,
- $C_{\mathrm{meas}}$ is the measured metabolite concentration,
- $f_{\mathrm{CSF}}$ is the CSF fraction within the voxel.

This calculation was performed independently for every participant using the measured CSF fractions before and after structural realignment. The resulting distributions of corrected concentrations were then summarized by their mean and standard deviation across participants below:

| **ROI** | **Before realignment** | **After realignment** | **Reduction in SD** |
|---|---|---|---|
| PFC | 1.207 ± 0.113 | 1.215 ± 0.111 | **1.5%** |
| Thalamus | 1.270 ± 0.288 | 1.155 ± 0.057 | **80.4%** |
| LSTG | 1.162 ± 0.109 | 1.157 ± 0.073 | **32.9%** |

Realignment substantially reduced the inter-subject variability of tissue-corrected metabolite estimates, particularly in the thalamus, where several subjects exhibited markedly elevated CSF fractions before realignment. These results indicate that realignment improves reproducibility of tissue-corrected metabolite quantification through reduction of variability arising from inconsistent anatomical sampling.
